# RADI (Reduced Alphabet Direct Information): Improving execution time for direct-coupling analysis

**DOI:** 10.1101/406603

**Authors:** Bernat Anton, Mireia Besalú, Oriol Fornes, Jaume Bonet, Gemma De las Cuevas, Narcís Fernández-Fuentes, Baldo Oliva

**Affiliations:** Structural Bioinformatics Lab (GRIB-IMIM), Department of Experimental and Health Science, University Pompeu Fabra, Barcelona 08005, Catalonia, Spain; Departament de Matemàtiques i Informàtica, Universitat de Barcelona, Catalonia, Spain; Centre for Molecular Medicine and Therapeutics, Department of Medical Genetics, BC Children’s Hospital Research Institute, University of British Columbia, 950 28th Ave W, Vancouver, BC V5Z 4H4, Canada; Laboratory of Protein Design & Immunoengineering, School of Engineering, Ecole Polytechnique Federale de Lausanne, Lausanne 1015, Vaud, Switzerland; Institut für Theoritische Physik, School of Mathematics, Computer Science and Physics, Universität Innsbruck. A-6020 Innsbruck, Austria; Institute of Biological, Environmental and Rural Sciences, Aberystwyth University, SY233EB Aberystwyth, United Kingdom; Department of Biosciences, U Science Tech, Universitat de Vic-Universitat Central de Catalunya, Vic 08500, Catalonia, Spain

## Abstract

**Motivation:** Direct-coupling analysis (DCA) for studying the coevolution of residues in proteins has been widely used to predict the three-dimensional structure of a protein from its sequence. Current algorithms for DCA, although efficient, have a high computational cost of determining Direct Information (DI) values for large proteins or domains. In this paper, we present RADI (Reduced Alphabet Direct Information), a variation of the original DCA algorithm that simplifies the computation of DI values by grouping physicochemically equivalent residues.

**Results:** We have compared the first top ranking 40 pairs of DI values and their closest paired contact in 3D. The ranking is also compared with results obtained using a similar but faster approach based on Mutual Information (MI). When we simplify the number of symbols used to describe a protein sequence to 9, RADI achieves similar results as the original DCA (i.e. with the classical alphabet of 21 symbols), while reducing the computation time around 30-fold on large proteins (with length around 1000 residues) and with higher accuracy than predictions based on MI. Interestingly, the simplification produced by grouping amino acids into only two groups (polar and non-polar) is still representative of the physicochemical nature that characterizes the protein structure, having a relevant and useful predictive value, while the computation time is reduced between 100 and 2500-fold.

**Availability:** RADI is available at https://github.com/structuralbioinformatics/RADI

**Supplementary information:** Supplementary data is available in the git repository.

## 1 Introduction

Protein structure is conserved through evolution, as protein function is structure-dependent (Lewis, et al., 2015). The reason for such conservation is due to energetically-favorable interactions between specific protein residues. Thus, there must be a certain degree of coevolution between the residues responsible for both the function and fold of all the members of a protein family (Schaarschmidt, et al., 2018). In the last decade, several authors developed the mean field approximation for direct-coupling analysis (DCA), either solving an inverse covariance matrix (Marks, et al., 2011) or using a pseudo-likelihood-based approach (Buchan and Jones, 2018; Ekeberg, et al., 2013) to compute direct information (DI) values (Giraud, et al., 1999) and detect causal correlated positions of the sequence. These correlations are reflected by co-evolution and potentially due to the spatial proximity of the residues in this position, thus helping to infer the contact map of a protein family (Morcos, et al., 2011). This has been used to improve protein models (Feinauer, et al., 2014; Michel, et al., 2014) or predict the structure of proteins (Hopf, et al., 2012; Ovchinnikov, et al., 2015). Here we propose a modification of the DCA algorithm for proteins that largely reduces its computational time, allowing the analysis of large proteins in reasonable time. Instead of analyzing every mutation in the protein, we ignore the mutations occurring within certain subsets of amino acids with equivalent physicochemical properties. We transform the sequences of a multiple-sequence alignment (MSA) of a protein family into a simplified alphabet of equivalent residues, and if the information of the alignment is still sufficient we calculate the top-ranking pairs of positions with high values of DI.

## 2 Methods

DI values are computed using a modification of the DCA algorithm, which we have named reduced alphabet DI (RADI). We denote by q the number of different symbols (i.e. alphabet) in the MSA.

### 2.1 Multiple-sequence alignments

MSAs are created using the script “buildmsa.py” included in the RADI Git repository. First, the script builds a profile of the query searching for similar sequences in the uniref50 database with MMseqs2 (Steinegger and Soding, 2017). Next, it uses the query profile to find more sequence relatives in the uniref100 database. Then, the script builds a MSA of the query and the identified sequences (up to 100,000) with FAMSA (Deorowicz, et al., 2016). Finally, it removes the columns of the MSA with insertions in the query. Note that MMseqs2 is executed with options “-s 7.5” and “--max-seq-id 1.0” for a more sensitive search.

### 2.2 Reduced alphabet

RADI simplifies the computation of DI values by transforming the original alphabet of q = 21 symbols (i.e. the 20 different amino acids plus the gap) into a reduced alphabet. For instance, using an alphabet of q = 21 (i.e. RA0) in RADI is equivalent to using the original DCA algorithm. We create three reduced alphabets (i.e. RA1, RA2, and RA3) by grouping amino acids based on different physicochemical properties (Table 1). We also define the number of effective sequences as the number of sufficiently different sequences (i.e. less than 80% of sequence identity) after removing the columns with more gaps than a threshold, which varies between 15% and 75% of the total number of sequences. Specifically, the threshold is set to the smallest percentage that would result in either 1,000 or the maximum number of effective sequences. Finally, we calculate the frequencies of symbols and weights them by the corresponding number of effective sequences at each position (Morcos, et al., 2011) to calculate the mutual or direct information, correcting the entropic effects with the Average Product Correlation (APC) (Dunn, et al., 2008).

**Table 1.**
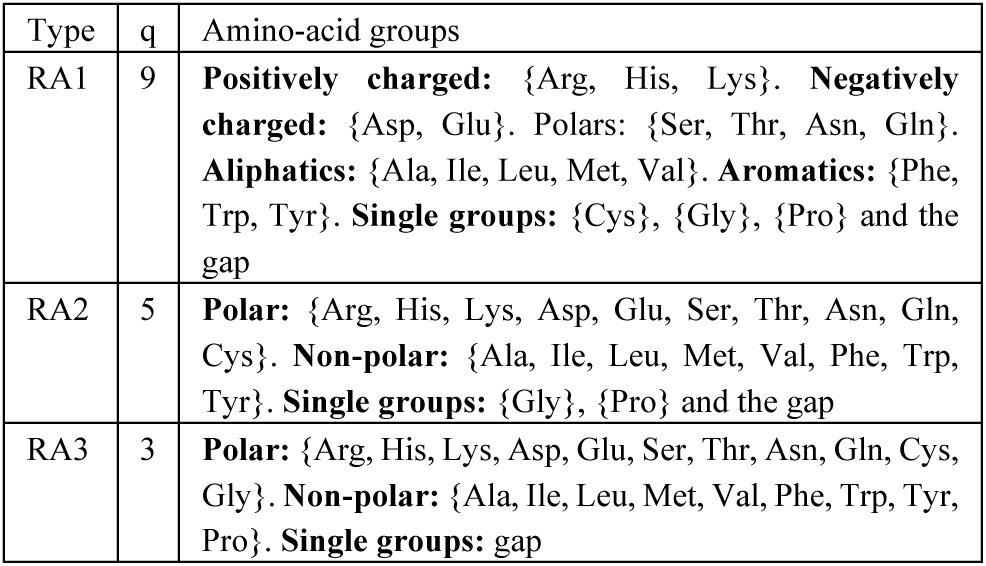
Classification of amino acids into groups for the three reduced alphabets (RA1 to RA3). The second column shows the number of *q* symbols for that reduced alphabet.

### 2.3 Analyses

Two residues are in contact if: 1) at least two atoms, one of each residue, are at a distance shorter than 5Å; or 2) the distance between their C[inline]atoms is shorter than 15Å; or 3) the distance between their C[inline]atoms is shorter than 8Å. We define the contact-map of a protein as the set of pairs of residues in contact. We only analyse the top DI pairs where the residues belong to two different secondary structures. We consider that two pairs, let them be (i,j) and (n,m), are equivalent if one of them is in the vicinity of the other defined by a [9×9] square (i.e. (i,j) is in the set of pairs [n-4,n+4]x[m-4,m+4]). Then, true positive contacts are top DI pairs equivalent to pairs of residues in the contact map. Top DI pairs using one of the RADI approaches (RA1, RA2, RA3) are similar to the original DCA method if they are equivalent to a pair predicted in RA0.

### 2.4 Hardware

To enable the benchmarking, RADI was tested on the queues of a cluster with the same CPU: 2 AMD Opteron 4226 hexacore of 2.9Ghz CPU with 64GB RAM.

## 3 Results

The modification of the alphabet results in different matrices of DI values. Nevertheless, we show that regardless of the alphabet, the top 40 pairs similarly hit equivalent residue-residue contacts with all alphabets, while reducing the number of symbols (*q*) greatly reduces the execution time.

### 3.1 Comparison of top DI pairs with respect the contact map

We compare both RADI and the original DCA algorithm on the same set of 509 different proteins from (Marks, et al., 2011), hereafter defined as benchmark. Protein sequences and three-dimensional structures are downloaded from the RCSB Protein Data Bank (PDB) (Rose, et al., 2017). As an example, we show one contact map for the molybdate binding protein (PDB code 1ATG), compared with the top 40 DI (and MI) values using the original DCA algorithm (*i.e.* RA0) and the reduced alphabets RA1 and RA3 (see figure 1A). For the whole benchmark (see details in supplementary data), we compare the distribution of the number of true positive contacts (Figure 1B). The averages of true positives across the 509 proteins vary between 29 (for RA3) to 35 (for RA0) and, although all distributions are significantly different, alphabets with RA0 and RA1 classifications are only slightly better than RA2 and RA3. Furthermore, the average number of similar top pairs of classifications RA1, RA2 and RA3 with respect to RA0 varies between 19 and 29, with more than 20 equivalent pairs between RA1 and RA0 for most proteins of the benchmark (Figure 1C).

**Figure 1.**
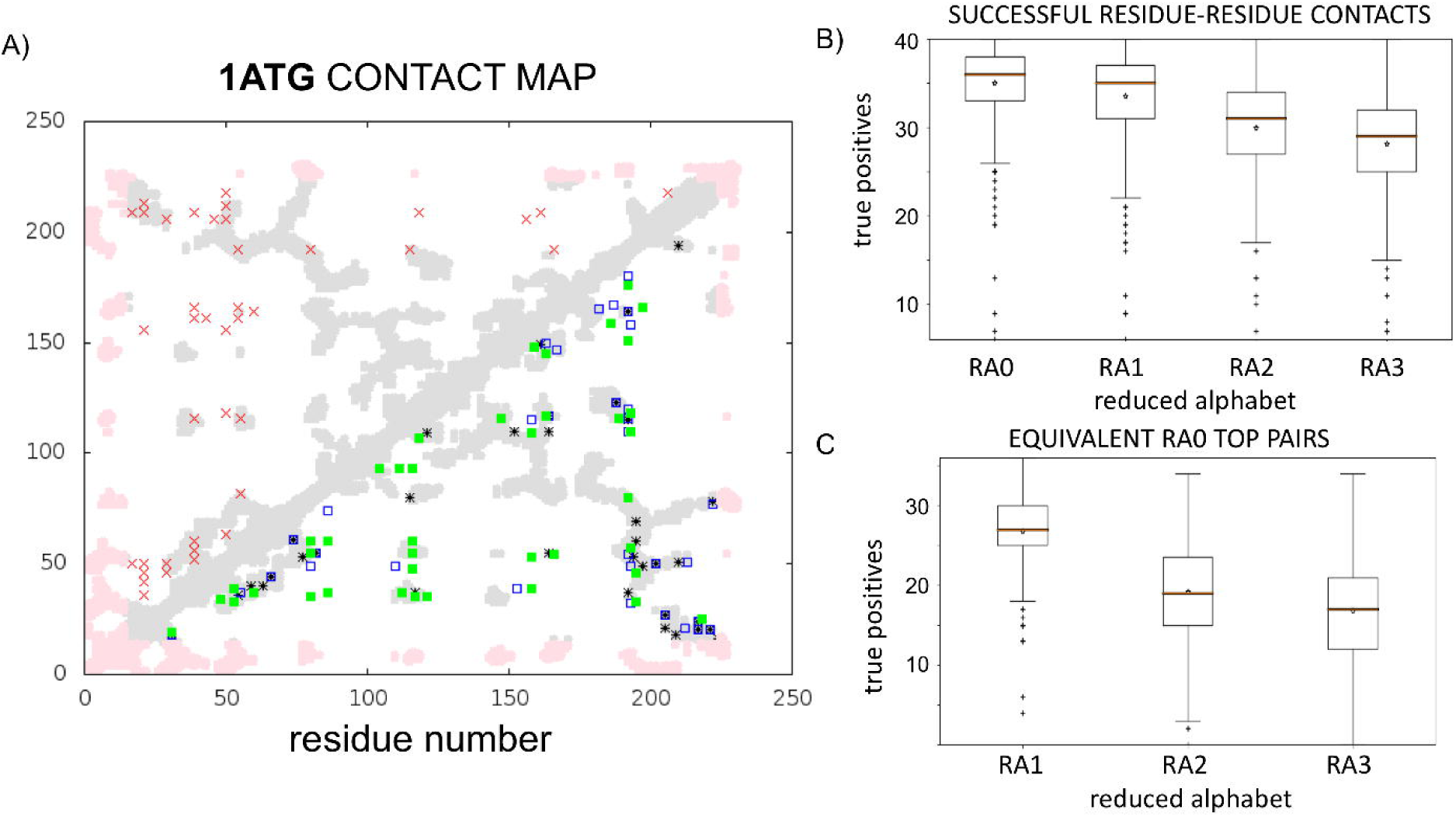
A) Example of residue-residue contact predictions for Molybdate binding protein (PDB code 1ATG). Real contacts are shown in grey (or pink if not sufficiently covered by the MSA). Red crosses show the 40 top pairs with higher MI values using RA0. Under the diagonal are shown the 40 top pairs with higher DI values using different amino-acid alphabets: RA0 (black stars); RA1 (unfilled blue squares); and RA3 (green squares). B) Distribution of the number of true positive contacts within the 40 top values of DI. C) Distribution of the number of residue-residue pairs in the 40 top DI values (for RA1, RA2 and RA3) equivalent to one of the top 40 DI values with RA0.

### 3.2 Improvement of execution time

The most computationally expensive step of DCA is the calculation of the pseudoinverse of a matrix, whose dimensions depend on the length of the protein (*L*) and the number of symbols (*q*) in the MSA alphabet. Reducing the alphabet from RA0 to RA1 speeds the computation time for a protein of *L* ≈ 900 by 32-fold, while the computation time when reducing the alphabet from RA0 to RA3 is ~2500-fold faster (Figure 2).

**Figure 2.**
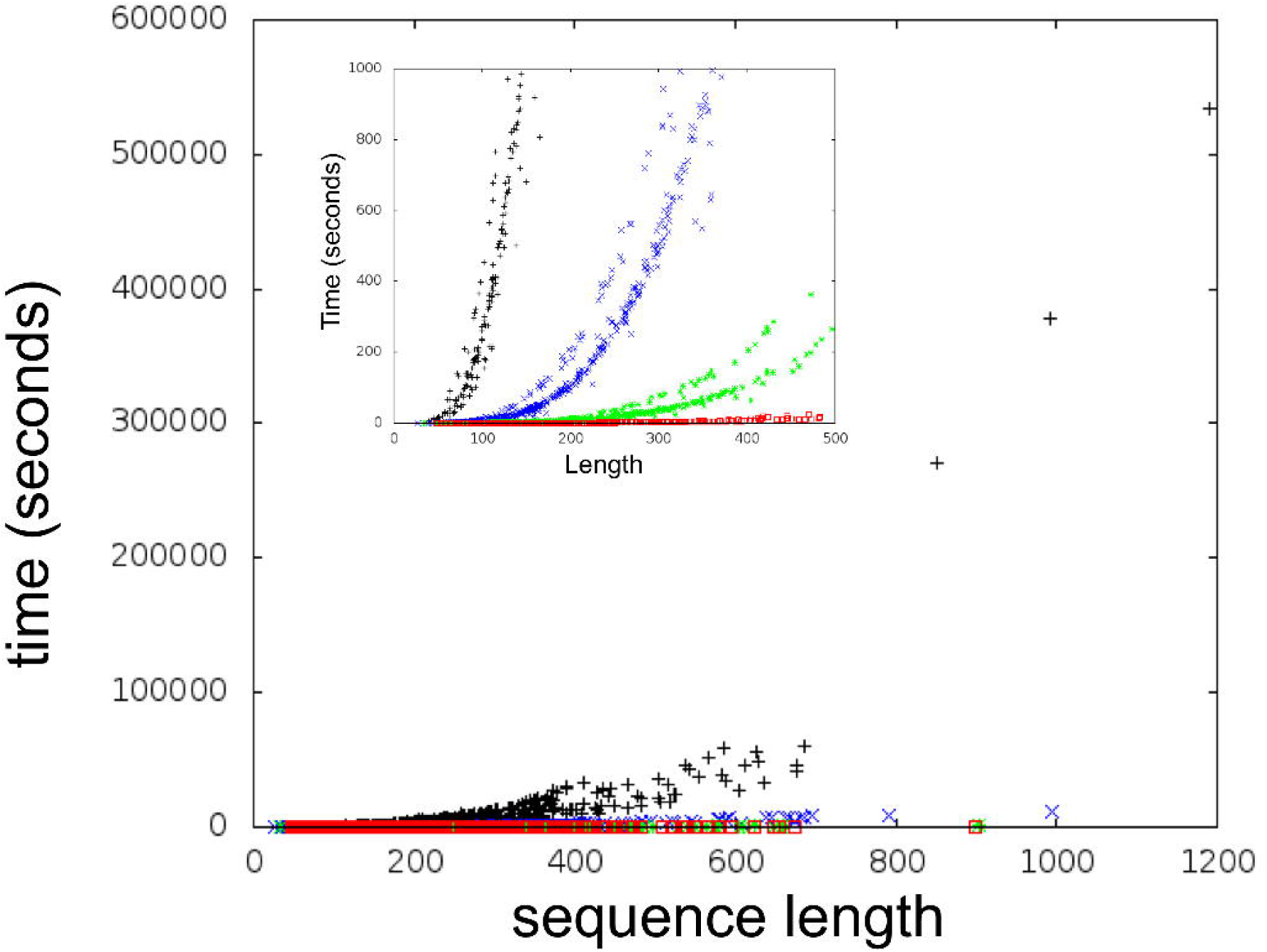
Computation times for different alphabets. Computation time of DI values as a function of the effective number of residues (length of the amino-acid sequence with sufficient sequences aligned in the MSA). DI values are calculated using all alphabets: RA0 (black); RA1 (blue); RA2 (green); and RA3 (red). The inner plot shows the behavior of the curves at the origin of the axes.

## 4 Conclusion

Regardless of the alphabet, top DI pairs can be similarly used for predicting the contact map of a protein structure. Consequently, we have to note two relevant conclusions. First, the time improvement in the computation of DI values enabled by RADI widens the applicability of DCA to proteins with more than 600 residues. For instance, for a protein of *L* = 600, the computation of DI values is largely reduced: from over 12 hours to merely 10 minutes when using the RA2 classification. And second, the simplification of the system produced by the reduction in the number of symbols is still representative of the physicochemical nature that characterizes the protein structure.

